# Reproducibility and replicability of rodent phenotyping in preclinical studies

**DOI:** 10.1101/079350

**Authors:** Neri Kafkafi, Joseph Agassi, Elissa J. Chesler, John C. Crabbe, Wim E. Crusio, David Eilam, Robert Gerlai, Ilan Golani, Alex Gomez-Marin, Ruth Heller, Fuad Iraqi, Iman Jaljuli, Natasha A. Karp, Hugh Morgan, George Nicholson, Donald W. Pfaff, S. Helene Richter, Philip B. Stark, Oliver Stiedl, Victoria Stodden, Lisa M. Tarantino, Valter Tucci, William Valdar, Robert W. Williams, Hanno Würbel, Yoav Benjamini

## Abstract

The scientific community is increasingly concerned with cases of published “discoveries” that are not replicated in further studies. The field of mouse behavioral phenotyping was one of the first to raise this concern, and to relate it to other complicated methodological issues: the complex interaction between genotype and environment; the definitions of behavioral constructs; and the use of the mouse as a model animal for human health and disease mechanisms. In January 2015, researchers from various disciplines including genetics, behavior genetics, neuroscience, ethology, statistics and bioinformatics gathered in Tel Aviv University to discuss these issues. The general consent presented here was that the issue is prevalent and of concern, and should be addressed at the statistical, methodological and policy levels, but is not so severe as to call into question the validity and the usefulness of model organisms as a whole. Well-organized community efforts, coupled with improved data and metadata sharing, were agreed by all to have a key role to play in identifying specific problems and promoting effective solutions. As replicability is related to validity and may also affect generalizability and translation of findings, the implications of the present discussion reach far beyond the issue of replicability of mouse phenotypes but may be highly relevant throughout biomedical research.

## Introduction

In recent years the scientific community has become increasingly concerned with cases of published “discoveries” that could not be replicated in subsequent studies, and sometimes could not even be reproduced in reanalysis of the original data. Such evidence is increasingly seen as a problem with the scientific method, questioning the credibility of science as a whole. Prominent institutions and journals, including the NIH, the NAS, *Science*, and *Nature*, have recently announced reconsideration of their policies due to this issue. However, there is still confusion and controversy regarding the severity of the problem, its causes, effective ways of addressing it, and what should be done about it, how, and by whom.

In the field of mouse phenotyping, the issue of replicability and reproducibility had been raised even before it became a widespread concern, and currently the NIH considers it to be especially prevalent in preclinical research. In mouse phenotyping the issue seems further tied to several other complicated methodological challenges, such as handling the potentially complex interaction between genotype and environment, defining and measuring proper behavioral constructs, and using the mouse as a model animal for human disease and disorder. In January 2015, researchers involved in the study of reproducibility and replicability gathered in Tel Aviv University to discuss these issues. These researchers come from various disciplines including genetics, behavior genetics, behavioral neuroscience, ethology, statistics, bioinformatics and database programming.

The present paper comprises of nine sections dedicated to nine central themes. In each section we attempt to present the general consent or at least a majority opinion, while more personal or controversial positions are attributed to researchers that support them, and links to meeting talks (available to the public in video clips through the Tel Aviv Replicability website) are referenced. All authors agree that this paper reflects the complexity of the replicability and reproducibility issues, even when restricted to a single area of research, yet it also points at practical ways to address some of these issues.

## 1. Reproducibility and replicability in general science: a crisis?

The ability to verify empirical findings wherever and whenever needed is commonly regarded as a required standard of modern experimental science. This standard was originally established in the 17th century, by Robert Boyle and other scientists of the Royal Society. These pioneers of experimental science regarded the ability to replicate results as an acid test differentiating science from one-time “miracles”. Their criterion for a scientific fact was (following a then common judicial dogma of two witnesses required for a valid testimony) something measured or observed in at least two independent studies (Agassi, 2013). In a case that may have been the first debate over the replicability of a scientific discovery, the Dutch scientist Christiaan Huygens noted a phenomenon related to vacuum in Amsterdam, and was invited to Boyle’s laboratory in London in order to replicate the experiment and show that the phenomenon was not idiosyncratic to his specific laboratory and equipment (Shapin and Schaffer, 1985). Ronald Fisher generalized the Royal Society criterion to more than two replications in his 1935 classic “The Design of Experiments”, writing: “we may say that a phenomenon is experimentally demonstrable when we know how to conduct an experiment which will rarely fail to give us statistically significant results” (Fisher, 1935). This quote illustrates how the common concept of statistical significance, already when it was first conceived, was closely tied with the concept of replicating experimental results. This concept served science well over the years, but recently non-replicated results have surfaced often.

In the field of mouse phenotyping, the problem has in fact always been present <u>^Crabbe^</u>, and was recognized before many other fields in the influential report by Crabbe and colleagues (1999). However, the issue is by no means unique to the field of mouse phenotyping. For instance, difficulties in replicating discoveries when dissecting the genetics of complex traits motivated the statistical threshold guidelines outlined by Lander and Kruglyak (1995).

Some notorious recent examples of poor credibility in general science include non-replicable methods of cancer prognosis (Potti et al. 2006, refuted by Baggerly and Coombes, 2009, and retracted), “Voodoo correlations” in brain imaging (Vul et al., 2009), “p-value hacking” (Simmons et al., 2011) and Excel coding errors that affected global economic policies (Pollin, 2014). A large community effort (Open Science Collaboration, 2015) recently attempted to replicate the findings of 100 papers in several leading psychology journals, and reported that 64% of the replications did not achieve statistical significance (but see Gilbert et al., 2016). A similar replication project in the field of cancer research (Errington et al., 2014) has not yet reported its results. The current situation is sometimes referred to as the “credibility crisis”, “replicability crisis” (e.g., Savalei & Dunn 2015), or “reproducibility crisis” (e.g., Peng, 2015) of recent science, and led prominent scientific journals and institutes to reconsider their policies (Landis et al., 2012; Nature Editorial, 2013; Collins and Tabak, 2014, who specifically mentioned preclinical studies as prone to reproducibility and replicability problems; McNutt, 2014; Alberts et al., 2015). Yet it is not clear what the new policies should be.

Ironically, there is currently no scientific consensus even over the name of the problem and the meaning of basic terms, confusing the discussion even further (Goodman et al., 2016). The terms replicable, reproducible, repeatable, confirmable, stable, generalizable, reviewable, auditable, verifable and validatable have all been used; even worse, in different disciplines and fields of science, these terms might have orthogonal or even contradictory meanings <u>^Stark^</u>. Following the now common term “Reproducible Research” in computer science (Diggle and Zeger, 2010; Stodden, 2010; 2013), a useful distinction was offered by Peng (2011, 2015) and Leek (Leek and Peng, 2015): “reproducibility” is concerned with reproducing, from the same original data, through analysis, the same results, figures and conclusions reported in the publication. “Replicability” is concerned with replicating results of another study, in a similar but not necessarily identical way, for example at a different time and/or in a different laboratory, to arrive at similar conclusions. We will use the above distinction in the remaining sections. However, note that other researchers recently suggested a similar distinction with the opposite terminology (Kennet and Shmueli, 2015).

Another categorization (Stodden, 2013) distinguishes between empirical reproducibility, computational reproducibility and statistical reproducibility. Stodden (2010, 2013) suggested that computational reproducibility is currently the most problematic. When viewing the objective of the scientific method as “rooting out error”, the deductive branch of mathematics (statistics included) has already developed its standards for mathematical proof, and the empirical branch (life sciences and mouse phenotyping included) has already developed its standards for hypothesis testing and method reporting. It is the third branch of computation-based research, that has yet to develop its own standards for reproducibility, which include data and code sharing (Stodden et al., 2013).

Ostensibly, science should not require trusting authority – it should be “show me”, not “trust me” <u>^Stark^</u>. Yet in reality, most scientific publications today amount to saying “trust me”. The typical scientific paper nowadays does not give access to the raw data, the code and other details needed to recheck the reported results – it basically says “I did all these carefully, you should trust my results”. A recent paper by David Soergel (2014) suggests that software errors (bugs) are not limited to a few high-profile cases that lead to retraction, and instead estimates that “most scientific results are probably wrong if the data passed through a computer”. In another estimation of the current state of science reproducibility, ThermoML, an open data journal in the field of thermodynamics, found that 20% of papers that otherwise would have been accepted contained serious errors (ThermoML, 2014). In summary, while we can strive for perfection the scientific process does not assure error-free results. What really blocks progress is the inability to detect and correct errors in time.

If a substantial percentage of published studies are not *reproducible* — i.e., if it is difficult to generate the figures, tables, and scientific conclusions starting from the data used in that study — it is unlikely that their results are *replicable*, i.e., that other researchers would be able to get similar numerical results and similar conclusions starting with data generated by a different researcher in a different laboratory. Within-study reproducibility seems a necessary but insufficient condition for across-study replicability of results. Ioannidis’ (2005) famous paper titled “Why most published research findings are false” highlighted the fact that the usual 0.05 significance testing increases the proportion of false discoveries among the discoveries made when testing many hypotheses. The combination of multiplicity and unadjusted testing can indeed be hazardous, as already argued by Soric (1987). In this case, emphasis on the use of 0.05 level testing encouraged superfluous solutions, such as the New Statistics movement (Cummings, 2014), which brands p-values as the source of the problem, and advocates replacing them with confidence intervals. In at least one psychology journal today, publishing p-values has been banned (Savalei and Dunn, 2015). However, in most cases the highlighted confidence intervals are still selected from many hypotheses, leading to the same issue of multiple comparisons (and even making it worse, since multiple comparisons corrections are less developed for confidence intervals than for p-values) <u>Benjamini</u>.

The alternative to discarding the p-values is to adjust them to cope with the multiplicity of hypotheses being tested or confidence intervals being made. The paper of Soric motivated the development of a formal approach to the False Discovery Rate (FDR) concept and the methods to control it (Benjamini and Hochberg, 1995). In the context of multiple comparisons it is easy to see why the “credibility crisis” in science became much worse in recent years: in the past, a typical phenotyping experiment would test a single measure of interest, or at most several measures. In recent years, however, automatized and computerized high-throughput strategies, such as the “batteries” (Brown et al., 2000) and “pipelines” of tests used for mouse phenotyping, frequently record ~10^2^ phenotypic measures, which is still far fewer than the 10^3^ –10^5^ variables measured in, e.g., typical GWAS studies. There is no way of reporting them all in a paper, so by necessity only a few are selected for highlighting. If the significant ones are selected as discoveries, the relevant error is expected to be the number of spuriously significant differences among the total number of significant differences. This equals the number of false discoveries among the total number of discoveries, namely the FDR. Recent attempts (Jager and Leek, 2014; Benjamini and Hechtlinger, 2013) to empirically estimate the “science-wise FDR” indicated a rate of 15–30%, which is considerably lower than Ioannidis’ warning of >50%, though also considerably higher than 5% (the commonly-used if arbitrary 0.05 significance level). These analyses also indicate that, once selective inference is accounted for, a 5% rate is indeed achievable. The Benjamini-Hochberg procedure of FDR is readily applicable to mouse phenotyping (Benjamini et al., 2001), and using it, especially in high-throughput multiple measure studies, should go a long way to decrease “discoveries” that do not replicate in later studies.

In summary, science is clearly reviewing its own failures, searching for the causes of the “crisis” and the ways to address them. The old concepts of experimental science emphasizing replicability and reproducibility are still correct in spirit, but seem to require updating of experimental, computational and statistical methodologies to cope with the increasing size and complexity of current experimental approaches. Pre-clinical research and mouse phenotyping are similar in this sense to other fields of science, but they encounter particular issues of their own.

## 2. Standardized mouse genotypes: replicated animal models?

The laboratory mouse is the main mammalian model animal for high-throughput genomic and behavior genetic research, and is used primarily to study the function of genes, a subject that is sometimes considered to be “the great challenge facing biologists today” (Collins et al., 2007). As an animal model, the laboratory mouse is used for pre-clinical development and testing of treatments for disease in humans, in genomic research and also in research of the Central Nervous System (CNS) and behavior, including studies by several participants in the meeting <u>^Pfaff,^</u> <u>^Tucci^</u>, <u>^Stiedl, Gerlai, Crusio, Golani^</u>. For obvious reasons, the reproducibility (within the same study) and replicability (across studies) of phenotyping in mice might have crucial implications for their relevance as animal models. Similar issues manifest in other model animals used for high-throughput phenotyping, such as the zebrafish <u>^Gerlai^</u>.

In the context of replicability, a main advantage of the mouse is our ability to standardize its genotypes, in the form of inbred strains or selected lines. Each inbred line consists of genetically-identical “clones”, which are in principle precisely replicated across studies. First generation (F1) hybrids of different inbred strains are genetically-identical as well (but not the subsequent segregating hybrid generations, due to recombination). Inbred strains and their F1 hybrids represent “genetic standardization” and collectively, sets of strains are referred to as “genetic reference panels”. They enable experimental designs that would be impossible using outbred animals or humans (monozygotic twins represent *n*=2, a sample size that imposes serious practical limitations). The BXD recombinant inbred lines, for example, can be thought of as “a cloned family” <u>^Williams^</u>, in which the C57BL/6J and DBA/2J genotypes are the parents, currently having about 150 reproducible “offspring” inbred lines, each representing a particular recombinant genotype between the parental genomes. Another such “cloned family” is the Collaborative Cross (The Complex Trait Consortium, 2004; Collaborative Cross Consortium, 2012), with eight “parents” and currently also about 150 “offspring” lines (Chesler et al., 2008; Iraqi, 2008; Morahan et al., 2008; Welsh et al., 2012, Iraqi et al., 2012). Quantitative Trait Locus (QTL) mapping methods (Complex Trait Consortium, 2003), a common analysis method discussed by several meeting participants ^<u>Chesler, Iraqi,</u>^ now routinely use the BXD and Collaborative Cross lines to localize quantitative phenotypes to segments of chromosomes within intervals of 0.5 to 2.5 Mb — often more than adequate to define and even clone causal sequence variants (Koutnikova et al., 2009; Houtkooper et al., 2013; Keeley et al., 2014). ^<u>Tucci</u>^

In parallel with the QTL approach, another method frequently employed is known as the “knock out” technology. It allows the targeted disruption of the gene of choice in the mouse (and more recently also in the rat). The result of the “knockout” is the generation of mutant mouse (or rat) lines in which a single gene is disabled, or mutated in a manner that would resemble a human genetic alteration. The goal of this technology has been the discovery of the effects of the mutation, which may aid our understanding of the biological functions of the targeted gene. That is, the knockout mouse has to be phenotyped. The International Mouse Phenotyping Consortium (IMPC, Beckers et al., 2009), a community effort for generating and phenotyping mice with targeted knockout mutations, discussed in the meeting by several IMPC researchers ^<u>Tucci, Karp, Nicholson & Morgan,</u>^ has a long-term goal to knock out most of the ~20,000 mouse genes, and phenotype them on the background of the C57BL/6 mouse genome, with the objective of annotating the functions of these genes.

The QTL and knockout approaches may be thought of as opposite strategies, respectively: “forward genetics”, which searches for loci within genes for a given phenotype, and “reverse genetics”, which searches for the phenotypic effects of a given genotype or genotypic manipulations. However, these two strategies are complementary and can increasingly be combined ^<u>Chesler, Williams</u>^ (Williams and Auwerx 2015; Wang et al., 2016). Understanding the phenotypic effects of mammalian genes is essentially the challenge of personalized medicine: if it is not possible to achieve in laboratory mice, where hundreds of individuals can be studied of each genotype, there is little hope to personalized medicine in humans <u>^Williams^</u>.

It is, however, important to recognize that genetic standardization in principle is not always standardization in practice. Non-replicable results sometimes reflect the naiveté of our expectations: they might result from heterogeneity in protocol, but also from a failure to recognize the importance of potentially subtle differences in genetic background ^<u>Valdar &</u> <u>Tarantino</u>^. For example, genetics might predispose some inbred strains to be more phenotypically variable. Highly homozygous genomes might have less capacity for "genetic buffering" against environmental variation, and some strains will be worse than others in _this respect_ <u>Valdar & Tarantino</u> (but see also Crusio, 2006).

Assuming such issues can be controlled, standardized mouse genotypes enable the replication of phenotyping experiments, and consequently studying the relation between genotype, environment and phenotype. This ability was a main motivation for conducting high-throughput mouse phenotyping replicability studies. If we can (with certain practical limitations and complications) replicate the genotypes, and if we manage (again, with certain practical limitations and complications) replicating the environment as well, shouldn’t we expect to replicate the resulting phenotypic effects? A major complication here is that the phenotype is a complex interaction between the genotype and the environment, and we almost never have the complete list of environmental factors and features that interact with the genotype or affect the studied subject. This complication is discussed in more detail in section 6.

## 3. Data sharing: how can it help with replicability?

Bioinformatics is a well-established discipline in the life sciences, traditionally concerned primarily with DNA, RNA and protein sequence data. The idea that phenotypic data are also worthy of storing, analyzing and reanalyzing (Gerlai, 2002) is not as well-established yet. Collaboration, community efforts, data sharing, and public phenotyping databases have an important role in today’s field of mouse phenotyping. Among many other utilities, they also offer unique opportunities for researching and controlling reproducibility and replicability. Many of the authors are personally involved with public databases and data sharing projects at different levels: from reanalyzing other researchers’ data to contributing their own data, and even constructing and maintaining public databases and community projects. Most presented in the meeting mouse phenotyping data collected across several laboratories, in some cases over long time periods, frequently through collaboration with researchers from other institutes and disciplines, and frequently contributing phenotyping data recorded in their own experiments to public databases and/or to reanalysis by other researchers ^<u>Benjamini, Chesler, Crabbe, Golani, Gomez-Marin, Heller, Iraqi</u>, Jaljule, <u>Kafkafi, Karp, Nicholson & Morgan, Tucci, Valdar & Tarantino, Williams, Würbel & Richter</u>^

A large data set used for analysis of replicability across laboratories (lab) was first presented in the meeting ^<u>Benjamini, Kafkafi,</u> Jaljule^, consisting of data from multiple databases and multi-lab studies contributed by several researchers, including Wolfer et al. (2004), Richter et al. (2011), Wahlsten and Crabbe (2006, downloaded from the Mouse Phenome Database) and knockout data downloaded from the IMPC database (Morgan et al., 2009; Koscielny et al., 2014). This dataset records results of individual animals (as opposed to just group means and standard deviations), amounting to one of the most extensive reanalysis of multi-lab studies, enabling estimation of the typical replicability in the field (section 4), as well as demonstrating the random lab model (see section 6) and GxL-adjustment (section 8) advocated for estimating replicability. GxL-adjustment explicitly relies on systematic data sharing as a proposed strategy for addressing replicability across laboratories in mouse phenotyping.

The Mouse Phenome Database (MPD, Grubb et al., 2014), a grant-funded research effort at The Jackson Laboratory, stores primarily phenotypes and protocol data, as contributed by researchers from over the world <u>^Chesler^</u>. It allows for trait correlation and examination of trait stability across strains, data sharing, dissemination and integration, facilitating the discovery of convergent evidence. Several among the meeting participants contributed their results to _the MPD_ <u>Crabbe,</u> <u>Chesler, Benjamini,</u> <u>Golani,</u> <u>Kafkafi,</u> <u>Valdar & Tarantino,</u> _and data from the MPD were_ used for several studies presented in the meeting. The GeneWeaver.org database (Baker et al., 2012), employs user-submitted or published gene sets from GWAS, QTL, microarray, text mining, gene co-expression, expert lists, curated annotations, and many other data sources drawn from major public data resources. It included (at the time of the meeting) ~80,000 gene sets from 9 species including humans and mice. GeneWeaver applies several algorithms to analyze the relations among these sets of genes. e.g., alcohol preference (Bubier et al., 2014, section 9).

GeneNetwork.org is a database that enables searching for ~4000 phenotypes from multiple studies in the BXD and in other recombinant inbred mouse families, as well as in other model organisms and in humans <u>^Williams^</u>. GeneNetwork employs a somewhat different strategy than the MPD in that it does not rely on researchers submitting their data. Instead the database operators have to extract the data from the scientific literature and integrate them into a uniform format. This requires a considerable effort, but also expands the range of studies and possible forms of analysis. GeneNetwork uses both routine and advanced statistical methods to extract, explore, and test relations among phenotypes and underlying genetic variation. It enables complex queries in real time, including very fast QTL mapping. It can also correlate any phenotype with all other phenotypes in the database across strain means, within or between studies. This makes it possible to explore the replicability of these phenotypes, even before relating them to the genotype. Any new phenotype can be correlated with any of the previous documented phenotypes across multiple strains. The increasing number of possible combinations, which grows exponentially with the rate of the added data, as noted “like good vintage: data sets get better with age” <u>^Williams^</u>.

The public database of the International Mouse Phenotyping Consortium (IMPC, Morgan et al., 2009, Koscielny et al., 2014) is intended to be “the first truly comprehensive functional catalogue of a mammalian genome”. The IMPC is a community effort to knock out ~20,000 genes and generate ~20,000 mutant mouse lines over the next 10 years, phenotype them using comprehensive and standardized high-throughput assays, and make them freely available to researchers over the world as animal models (De Angelis et al., 2015). At the time of the meeting the IMPC included ten “centers” – institutes over the world performing high-throughput phenotyping of mice, over the same genetic background of C57BL/6. Although most lines were tested only in one center, a few mutant lines and their C57BL/6 controls were tested across 3 and even 4 centers, and even more overlap between centers currently accumulates, enabling an extensive study of replicability across laboratories. The IMPC has made an effort to standardize phenotyping assay protocols across centers. In certain pipelines it may record ~270 phenotypic measures per mouse ^<u>Nicholson & Morgan, Karp.</u>^ Despite the standardization there is still workflow variation among centers, as a results of local factors such as different policies and colony size, for example, mice from the same litter are typically assayed on the same day, and some centers have concurrent controls while others regularly sample controls. Data from the IMPC database are currently being used for several studies of replicability _<u>Karp, Nicholson & Morgan, Benjamini, Kafkafi,</u>_ Jaljule.

## 4. Replicability issues in mouse phenotyping – how serious are they really?

This seemingly simple empirical question is not simple at all to answer, for several reasons. Only a few attempts at systematic analysis have been conducted across several studies and/or laboratories with the objective of estimating replicability in a quantitative way (see also sections 3 and 8). Furthermore, there is no consensus even over the correct ways to analyze and estimate replicability (Open Science Collaboration, 2015; Gilbert et al., 2016, see also section 6). Generally, most of the participants in the meeting seem to agree that there is indeed a real and serious problem of reproducibility and replicability in the field of mouse phenotyping. However, some specific phenotyping results appear highly replicable, especially when having large genotype effect sizes.

Crabbe et al. (1999) conducted the famous experiment that brought up the replicability issue in the field of mouse phenotyping, and preceded the current “credibility crisis” in general science (Ioannidis, 2005). This experiment compared five inbred strains, one F1 hybrid and one knockout line and its inbred background strain across three laboratories, standardizing factors including equipment, protocols, and husbandry at a much higher level than is common in the field. This study found, in most of the eight standard phenotypic measures, significant effects of the laboratory and/or interaction between laboratory and genotype, and therefore made the provocative conclusion: “experiments characterizing mutants may yield results that are idiosyncratic to a particular laboratory”. Additional results were published in another study across laboratories and across several decades of phenotyping (Wahlsten et al., 2006). On the other hand, several genotype differences appeared replicable, especially when genotype effect sizes were large.

Generally, John Crabbe ^<u>Crabbe</u>^ estimated that the issue has been exaggerated, and that the situation is actually not worse than that of many other fields of science. At the time a response paper (Pfaff et al., 2001) presented several effects of mutations in mice that were replicated. However, this does not ensure that any mutation effect discovered in a single laboratory, even a large one, will replicate in other laboratories.

In another study looking at nociception phenotyping, about 42% of the variance was found to be associated with the experimenter, and many other sources of laboratory
environmental variation were found to influence phenotype alone and in sex and genotype interactions (Chesler et al., 2002). Similar effects were found for many other behavioral phenotypes (Valdar et al., 2006). In QTL analysis using lines of the Collaborative Cross, different cohorts often produce different results ^<u>Iraqi</u>^, seemingly affected by factors such as season, time of testing in the circadian phase and even geographic altitude.

A common way to visualize the replicability across two experiments from different studies, or even different laboratories, is a correlation plot of the genotype means, e.g., in Wahlsten et al. (2006). Several speakers in the meeting presented such plots comparing laboratories and studies ^<u>Chesler,</u> <u>Crabbe,</u> <u>Valdar & Tarantino</u>, <u>Tucci</u>^, and both the MPD and the GeneNetwork software (see section 3) generates them by request, and even runs a fast search in its database for phenotypes that correlate with any given phenotype across strains <u>^Williams^</u>. Such plots frequently indicate considerable correlation (positive or negative) between strain means across studies, indicating some replicability, although there is no clear criterion for how much correlation indicates sufficient replicability.

A recent heterogenization experiment (Richter et al., 2011,) was orchestrated across six laboratories, more than in any multi-lab experiment in the field of mouse phenotyping. It concluded that these laboratories, while still much fewer than the number of all potential mouse phenotyping laboratories over the world, already contribute a large component of variation, apparently considerably larger than the variation introduced by systematic heterogenization of two factors (test age and cage enrichment). This study therefore concluded that “differences between labs are considerable and unavoidable” (this study and the heterogenization approach are discussed in more detail in section 7)

There are many potential confounds in the study of genetically modified mice that are difficult to control (Schellink et al., 2010) and they are likely to differ across laboratories and studies. Studies utilizing phenotyping data from several knockout lines and associated controls across research centers of the International Mouse Phenotyping Consortium (IMPC) were presented in the meeting ^<u>Karp,</u> <u>Nicholson & Morgan</u>^. These studies found at each phenotyping institute that test day was a considerable source of variation and encompassed multiple variance sources (e.g. human operator, litter, cage, reagents etc., see also Karp et al., 2014). Spreading mutants across time functions as a form of heterogenization ^<u>Karp, Nicholson & Morgan</u>^. It is not clear to what extent a multi-batch workflow captures the interaction of genotype with the laboratory, which is a different effect. Litter effects in the IMPC phenotyping data, if not corrected, might mask genotype effects in the (non-littermate) wild-type vs. mutant t-test ^<u>Nicholson & Morgan</u>^.

In a large dataset comprised of multiple previous studies, each including several genotypes measured across several laboratories (see section 3), cases were demonstrated that may be termed “opposite significant”: i.e., one genotype is actually significantly higher in one laboratory while significantly lower in another laboratory <u>^Kafkafi^</u>. In other words, each of these laboratories by itself would have reported the opposite discovery. Opposite significant cases are not rare: examples were found in most of the multi-lab datasets in the study, although as expected they are more common in datasets performed over a larger number of laboratories ^<u>Kafkafi</u> <u>Würbel & Richter</u>^. However, using the random lab model criterion for a replicable genotype effect (see section 6), in most multi-lab datasets (specifically all 8 but one) the majority of genotype effects were replicable <u>^Kafkafi^</u>. In these same datasets, the proportion of “non-replicable positives”, i.e., genotype differences that were found significant within a single lab (using the typical t-test at the level of α = 0.05) but did not replicate across all laboratories (using the random lab model) ranged between 20% and 40% ^Jaljule^. This result can be regarded as an estimation of the proportion of non-replicable “discoveries” in single-lab studies in the field. It could be argued to be slightly lower, because the above effect was calculated using the same results for both the single-lab and multi-lab estimation, but it could also be argued to be is higher, since the general standardization level in the field is probably lower than the standardization level in the multi-lab studies used to derive the above proportion.

Most speakers seem to agree that the proportion of non-replicable results in the field is considerably higher than the sometimes assumed (or hoped) 5%, yet they also believe that this proportion is not so high as to make the whole field worthless, and can be improved using several approaches (see sections 7, 8 and 9).

## 5. Replicability of behavior: a special case?

The recent concern in the scientific community regarding replicability and reproducibility of experimental results is by no means limited to behavioral studies, and Ioannidis’ (2005) famous claim that “most published scientific results are false” does not single them out. While psychology is frequently mentioned as a field that might suffer from a high percentage of non-replicable discoveries (Asendorpf et al., 2013; Open Science Collaboration, 2015), so are other fields, such as pre-clinical and clinical pharmacology, epidemiology, physiology, brain imaging and Genome Wide Association Studies (GWAS). Issues with replicability of a non-behavioral phenotype in mice – susceptibility to infection – were also presented in the Tel Aviv meeting <u>^Iraqi^</u>. However, several speakers <u>^Crusio, Eilam, Gerlai, Golani, Gomez-Marin, Pfaff, Stiedl^</u> noted that behavior is methodologically problematic to understand and measure (Gomez-Marin et al., 2014), and this probably hurts its replicability as well. This section follows their discussion as to how and why we should, highlighting methodological difficulties of measuring behavior and offering solutions.

Individual differences are common in behavior, increasing within-group noise and thus might be suspected to impede replicability. The two-compartment DualCage setup (Hager et al., 2014), while sensitive enough to differentiate the behavior of the two closely-related mouse substrains C57BL/6J and C57BL/6N, also revealed large inter-individual differences with some mice showing Post-Traumatic Stress Syndrome (PTSD)-like persistent avoidance performance. The generally short test duration of many high-throughput assays may imply lower variability than that detectable if animals are offered deliberate choice(s) over long time scales. In C57BL/6N mice, strong reinforcement resulted in two performance subgroups with either low or maximum avoidance, indicating weakened or PTSD-like memory, respectively. Performance differences in cognitive tests can emerge due to the differential impact of specific and unspecific stressors and emotional (anxiety) differences <u>^Stiedl^</u>. This particularly applies in complex tasks involving higher cortical functions thereby following the arousal-performance relation of the Yerkes-Dodson law (reviewed by Diamond et al., 2007) that has been known for more than 100 years. In contrast, other behavioral measures such as locomotor activity are highly correlated over successive days in the DualCage indicating high stability and therefore high replicability. In humans each motor task includes components necessary for its completion, but also “idiosyncratic acts” that are particular to the individual and instance. Idiosyncratic acts in the population are typically a considerable proportion of the total number of acts, and they appear to have important cognitive function (Eilam, 2014).

An interesting empirical question is whether behavioral phenotypes are less replicable than physiological phenotypes. A general consensus seems to be that behavioral phenotypes need not be less replicable. In a study across several laboratories and many decades, Wahlsten et al. (2006) showed that some behavioral phenotypes (including locomotor activity) were as replicable as classic anatomical phenotypes such as brain size, yet other behavioral phenotypes were considerably less replicable. Proekt et al. (2012) demonstrated that motor activity in home cages can be highly reliable, and as much as physical variables in exact mathematical models, providing some conditions were met <u>^Pfaff^</u>. Replicability problems often arise because of basic methodological and statistical issues. For example, in the analysis of the IMPC phenotyping data, issues with temporal fluctuations in control mice were observed in both behavioral and physiological assays <u>^Karp^</u>.

However, basic methodological issues with behavior research might negatively affect its replicability ^<u>Crabbe, Gerlai, Würbel & Richter</u>^. Especially, behavioral measurements might dramatically change as a result of unforeseen and unknown environmental effects, which may reduce statistical power and replicability. Such issues are not special to the mouse, and were also illustrated in zebrafish <u>^Gerlai^</u>. They might be worse in the high-throughput procedures used in phenotyping, which are typically specialized for efficiency and ease of measurement. Several speakers stressed that automation should be developed only on the basis of careful prior observation and thorough understanding of the behavior ^<u>Gerlai,</u>^ <u>Golani, Gomez-Marin, Pfaff</u>.

Many common behavioral constructs, such as “anxiety” and “behavioral despair”, are not sophisticated nor well validated, and understanding of what it is that assays for these constructs measure is insufficient <u>^Crusio^</u>. Standard and commonly-used behavioral tests are not immune to these issues. For example, the Porsolt Forced Swim Test and the Tail Suspension Test are both considered as measuring “behavioral despair” using a similar variable: the latency to stop trying to escape from an unpleasant situation; yet some mice treated with a random stress procedure reacted oppositely in these two tests <u>^Crusio^</u>. While researchers assume that these tests measure a similar construct, the mice apparently disagree. In a similar example, the Morris Water Maze is frequently considered to be a “gold-standard” for memory testing in rodents, yet in mice, factor analysis revealed that most of the variance is explained by behaviors such as wall hugging (thigmotaxis) and passive floating, and only about 13% by memory (Wolfer et al., 1998). Such methodological problems may negatively affect replicability as well.

Addressing such problems is critical when deciding how to share and integrate behavioral data, for example, when constructing “vocabularies” and “ontologies” used for querying and searching phenotyping databases <u>^Chesler^</u>. Semantics raises many challenging questions regarding how behavior should be described, including: (a) How are behaviors hierarchically related to one another? (b) How should they be organized? (c) Should they be labeled by the supposed meaning of the assay, or only by what can be observed? For example, “immobility” is an objective description of motor behavior free of the context of ascribed behavioral meaning. Mice are immobile in a variety of apparati for a variety of reasons and in response to treatments. Should an “immobility” label be used rather than labels such as “anxiety”, “fear,” “learning” and “behavioral despair?

Other speakers also agreed that the relevant variables affecting behavior are frequently not as well-understood as those of physiology, anatomy or genetics ^<u>Benjamini, Golani, Gomez-</u> <u>Marin, Kafkafi, Stiedl</u>^. On the other hand it is possible to consider replicability itself as the gold standard, and use behavioral data from several laboratories in order to explicitly redesign behavioral constructs for increased replicability <u>^Golani^</u>. In this framework, the issue of replicability in behavioral results is turned into an asset rather than a liability – it enables researchers to methodically improve the definition of key behavioral constructs by using the statistical level of replicability as a benchmark, filtering out behavioral results that are not replicable.

This issue is closely connected with the issues of genotype-environment interaction (see section 6) and validity of behavior (see section 9).

## 6. Genotype-Environment Interaction – how should it be handled?

A problem inherent to the field of phenotyping is that the final phenotype depends not only on the genotype and the environment, but also on an interaction between the genotype and environment (commonly abbreviated GxE). Furthermore, the effect of the environment on animals is cumulative, with phenotypic measures often depending on historic development and experiences of the animal. For example, in cross-fostering experiments of mouse inbred strains, raising BALB/cByJ pups by a C57BL/6ByJ dam reduced excessive stress-elicited hypothalamic-pituitary-adrenal (HPA) activity and behavioral impairments, but no effect was found in the opposite case of C57BL/6ByJ pups raised by a BALB/cByJ dam (Anisman et al., 1998). These interactions may take place over the many levels of RNA, protein, cells, tissues, whole-organisms and ontogenetic development. In the case of brain and behavioral phenotypes, there are the additional levels of neurons, CNS organization and activity, as well as their complex interaction with the environment. The physicist PW Anderson was quoted in the meeting <u>^Williams^</u> (1972): “surely there are more levels of organization between human ethology and DNA than there are between DNA and quantum electrodynamics, and each level can require a whole new conceptual structure”. This understanding is commonly regarded in current life sciences to be the answer to the old “Nature vs Nurture” debate, and the basis of the “system biology” and “whole-animal research” approaches, and is closely connected with the ecological concepts of phenotypic plasticity and reaction norms (Lewontin, 1974; Wahlsten, 1990; Pigliucci, 2001; Voelkl and Würbel, 2016) as well as the psychological concept of G-E correlations (Homberg et al., 2016).

Empirically, this biological interaction does not necessarily have to result in large statistical interaction between genotype and environment, but in many cases it does. The most obvious case of statistical GxE occurs when a certain genotype (e.g., a knockout inbred line) scores a higher phenotypic mean than another genotype (e.g., the wild-type inbred strain) in certain environmental conditions, yet lower in other environmental conditions (Fig. 1 left). Typical examples of different environmental conditions may be different laboratories, different test days at the same laboratory, or even different laboratory technicians. The sources of interaction are frequently unknown, multiple and very difficult to control. An intuitive hypothetical illustration would be a test whether Dobermans are more aggressive than Poodles <u>^Golani^</u>. While in one laboratory this indeed may be the result of a standardized aggression test, in another laboratory the lab technician might unknowingly wear a perfume that annoys only the Poodles, leading to an opposite result that might prove very difficult to “debug”. Such opposite results are quite common in actual mouse phenotyping ^<u>Benjamini,</u> <u>Kafkafi, Würbel & Richter</u>^ as well as in other model animals such as the zebrafish <u>^Gerlai^</u>. They are actually more impressive than the hypothetical dog breed example, since C57BL/6 mice, unlike Dobermans, are (near perfectly) genetically-identical. A large interaction effect is usually considered the mark of a true non-replicable genotype effect <u>^Kafkafi^</u>. Note that an environment effect alone is not a serious hindrance to replicability since, by definition, it affects all genotypes to the same amount, and therefore can be controlled by having measurements on control animals (e.g., a reference genotype). An interaction effect, in contrast, cannot be corrected this way because it is by definition unique to the specific combination of both genotype and environment.

**Figure. 1:**
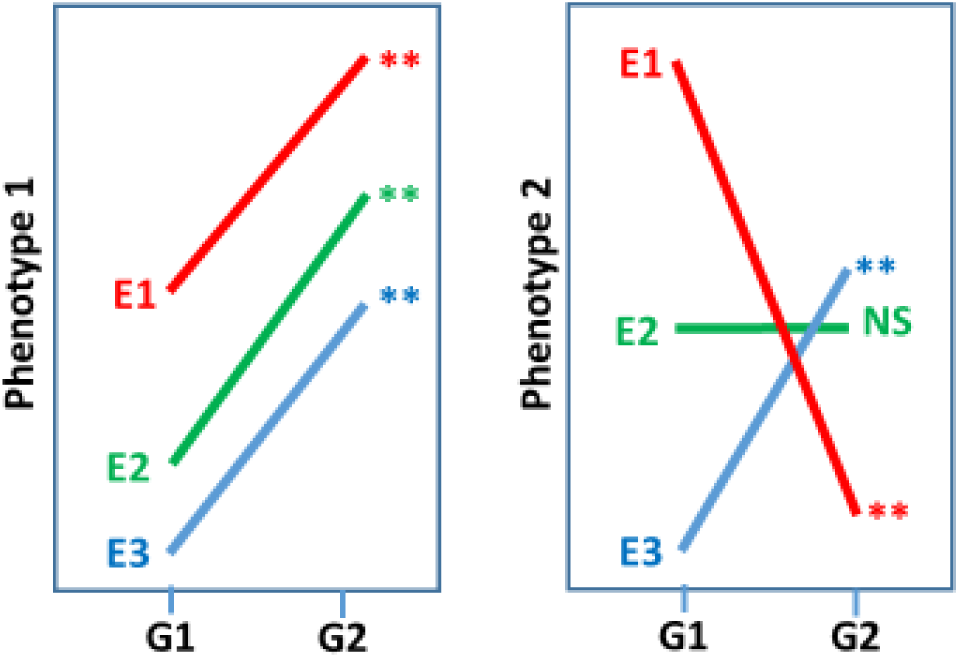
Comparing two genotypes G1 and G2, using two phenotypic measures 1 and 2, in three environments E1, E2 and E3. In the case of phenotype 1 (left) there is almost no interaction between genotype and environment (GxE). Note that the environment effect is large, but since it affects both genotypes in the same way it can be controlled using the same genotype as a reference for all other genotypes within the same environment. In the case of Phenotype 2 (right), there is a strong GxE effect, to the point that in E1, G1 is significantly larger than G2, while in E3, G1 is significantly smaller than G2 (“opposite significant”). In this case an issue with replicability ensues, since the genotype effect is idiosyncratic to the specific combination of genotype and environment.

What can and should be done about the statistical GxE interaction? This depends on the research question <u>^Benjamini^</u>. In many cases the source of the interaction, once recognized, might be itself of interest, and lead to uncovering an important biological or behavioral mechanism. However, when testing the very common type of hypothesis suggesting that a certain genotype has a certain phenotypic effect, the interaction is at least a confounding factor (Fig. 1B) that must be taken into consideration and handled, and is even considered by some to be a fundamental property of living organisms. As illustrated and discussed in the meeting, carful observation of the animals’ response to the rearing conditions and/or experimental setup may sometimes locate and eliminate the cause of some of the interaction ^<u>Crabbe,</u> <u>Crusio,</u> <u>Gerlai, Golani, Tucci</u>^. Moreover, certain phenotypic measures might be much less sensitive to GxE than other measures, especially if they are more robust to environmental disturbances and more faithfully represent the true state of the animal (Wahlsten et al., 2003; Benjamini et al., 2010). A systematic way of decreasing the interaction was demonstrated by explicitly changing and improving the measure used for phenotyping <u>^Golani^</u>.

In many cases a certain portion of the statistical interaction effect does not disappear even after carefully redesigning the experiment or improving the analysis, and remains large and significant. Large GxE interaction effects may still be highly replicable if they depend on well-known environmental condition that can be equated (such as the dependence of body size in drosophila strains on temperature) but often they do not. In such cases the common statistical approach in the field brands the genotype effect as non-replicable, being idiosyncratic to unknown sources and conditions. However, according to the newly developed ”Random Lab Model” (Kafkafi et al, 2005; and see section 8), such a genotype effect may still be demonstrated as replicable, providing it is large enough to be significant even over the background of the large interaction. The Random Lab Model treats the genotype as a fixed factor that can be precisely standardized and replicated, but models the environment with random variables. This approach gives up on the unrealistic hope of precisely standardized and replicated laboratories, and instead models them as randomly sampled out of the population of all phenotyping laboratories. The immediate implication is that the interaction of the genotype with the laboratory (GxL) has a similar role to that of the individual animal noise (within-group effect).

Similar to the individual animal noise, it should be decreased as much as possible, but in real life it would never disappear completely. Instead the model adds it to the within-group variability as the yardstick against which the genotype effect is compared. This generates a higher benchmark for showing a significant genotype effect – the price paid for ensuring that this effect is likely to remain significant if tested in another laboratory.

It is rarely appreciated that the most common criterion in the field for assessing replicability across several laboratories – the significance of the GxL interaction effect in the traditional fixed model ANOVA – often results in misleading (Kafkafi et al., 2005) and even paradoxical conclusions <u>^Kafkafi^</u>. Perhaps the worst is that using low-quality and noisy measurement may render the interaction non-significant. Alternatively, the same consequence can be “achieved” by using samples that are too small. In both cases the a semblance of replicability is created, The reason is that the standard fixed ANOVA has lower intrinsic power to detect interaction effects than to detect the main effects (Wahlsten, 2006), and thus any degradation of power is likely to eliminate GxL significance before it eliminates the genotype significance. This seeming paradox can be resolved by using the Random Lab Model instead of fixed model ANOVA. With this model, the criterion for a true genotype difference and the criterion for a replicable genotype difference are one and the same – the significance of the genotype effect. It is therefore impossible to “improve” replicability by degrading the power (Kafkafi et al., 2005).

Replicability issues in the same laboratory across time is a similar problem arising as a result of “workflow” – the timing of individual mouse testing, either knockout mutants or background controls <u>^Karp^</u>. In IMPC centers, each mouse passes through a phenotyping “pipeline” – a series of phenotypic tests in a predetermined order and defined age of the mouse. Due to fecundity and fertility problems, there are typically multiple small batches of knockouts with different birth dates and therefore testing dates, and the control mice (which are typically much larger in number) might not be concurrent. Moreover, depending on institutional resources and throughput, different institutes can have different workflow. Karp et al. (2014) preferred moving to a design and analysis which embraces this variation across time, rather than changing to a highly standardized design. She proposed a mixed model in which time (batch) is a random variable.

Handling GxE interaction of various kinds thus depends on the objective and context of research. GxE can be understood and decreased by carful observation of the animals, and by redesigning housing and testing conditions. Especially when testing a hypothesis of a genotype effect, ignoring or mishandling potential GxE is likely to result in replicability issues and other severe methodological issues.

## 7. Standardization and heterogenization: why and when should they be used?

When discoveries of preclinical studies fail to replicate despite the near-perfect standardization of the genotypes, there is a natural tendency to assume that the problem is the lack of standardization of housing and testing conditions. The standardization challenge seeks to document the important properties of the environment and then keep them constant. A commonly held ideal is that every animal will be treated identically, so there are no sources of variation other than the controlled variable used as experimental treatment. This common dogma is usually invoked in the context of the “internal validity” within one study in one laboratory. In this role standardization is seen as means to minimize the variability of results, avoiding bias by unwanted sources of variation, and increase sensitivity and precision. It is typically assumed that standardization lowers the noise level, therefore increases signal-to-noise ratio and increases the statistical power to show differences between the experimental and control groups, thus increasing significance and decreasing the number of animals required to show a given effect size and significance.

Several participants in the meeting invested considerable effort devising behavioral assays in which the animals are tested for a long time (24 hours and more) in a home cage, sometimes with additional space to explore, with minimal or no contact with a human experimenter, but potentially with computer-controlled experimental manipulations and feedback to increase standardization. Proekt et al. (2001) developed a high-throughput assay including multiple computer-controlled units, in which the mice are completely isolated from outside sound and vibration, and require human experimenters touch them only once per week. Tactile, olfactory and vestibular stimuli can be programmed, and the animal movement is tracked using infrared beams <u>^Pfaff^</u>. Fonio et al. (2009) video-tracked mice (as well as fruit flies) in assays comprised of a small home cage connected through a narrow doorway with a much larger arena, which the animals gradually and systematically inhabit over several hours to several days, of their own volition with no apparent incentive other than exploration <u>^Golani^</u>. Tucci’s laboratory at IIT(Maggi et al., 2014) developed automated home-cage testing (<u>www.phenoscale.org</u>), consisting of computer-controlled holes and hoppers, in which circadian rhythms, sleep-wake and related cognitive processes can be automatically recorded and studied for many days <u>^Tucci^</u>. Moreover, Tucci’s team has developed user-friendly software platforms that can handle raw data and make them available to the community to improve data sharing and to coordinatel multiple testing across different laboratories. Hager et al. (2014) developed a two-compartment home cage-based assay with video-tracking to monitor fear learning, avoidance and risk assessment over two days without human interference. Here individual variability in exploring a test compartment was detectable in the absence of the experimenter (see chapter 4) as a potentially confounding factor, indicating that the assumption that standardization may help lower variation may not apply to all behavioral measures (see chapter 5) <u>^Stiedl^</u>.

Standardization is employed for another objective: increasing reproducibility across replicates of an experiment, either across time within the lab or across multiple labs. Crabbe et al. (1999) made an exceptional effort to standardize their phenotyping experiment across three different laboratories <u>^Crabbe,^</u> and the EUMORPHIA project standardized the IMPC pipelines of tests across the IMPC centers <u>^Tucci^</u>. Both reported that careful improvement of conditions and testing improved replicability, yet both reported issues of replicability despite standardization.

Richter et al. (2009, 2010, 2011) maintain that the idea to improve replicability through standardization is based on the true finding that experimental results can differ depending on environmental conditions (i.e., phenotypic plasticity, Lewontin, 1974; Wahlsten, 1990; Pigliucci, 2001; Voelkl and Würbel, 2016), and on the false belief that these conditions are fully known so standardization will ‘spirit away’ such differences between experimental conditions, which they refer to as “the standardization fallacy” ^<u>Würbel & Richter</u>^. On the contrary, they proposed that “heterogenization” – systematic variation of conditions – may improve reproducibility and attenuate spurious results. The rationale is that different laboratories will always standardize to different local conditions, because many lab-specific factors are either unknown or cannot realistically be standardized, such as personnel. Consequently, the results might be valid only for these narrow conditions, and may not necessarily generalize to the conditions of other laboratories. In the last of several proof-of-concept studies, Richter et al., (2011) ran heterogenized and standardized batches in each of six laboratories. In this study, heterogenization of study populations through systematically varying animal age and cage enrichment did indeed improve replicability, but only by a very small extent. It appears that this simple form of heterogenization, introduced only a fraction of the variation that existed among the six laboratories. It is also notable that standardization may be a possible reason why preclinical studies often find drug efficacy while phase 2 or phase 3 human clinical trials of the same drug fail. Human populations are variable, genetically and environmentally, while animal populations are often genetically highly homogeneous and are tested under strict environmental control. These discrepancies have been discussed and the question of how to resolve them has debated in the pharmaceutical industry and academia alike.

Several speakers in the meeting were supportive of the heterogenization concept, and (perhaps surprisingly in light of the importance usually prescribed to standardization) no one objected to it outright. Notably, John Crabbe argued that strict “universal” standardization of laboratory and test environment is unfeasible <u>^Crabbe^</u>. In his opinion widespread adoption of fewer protocols, apparatuses and test environments would diminish, not enrich, understanding of the assayed behavior. Any attempt to repeat an experiment can never perfectly replicate the original conditions, but this is probably a good thing since it will increase generalizability of findings. Similarly, IMPC researchers in the meeting, separately analyzing mouse phenotyping data from the IMPC, noted that spreading mutants across time can be regarded as environmental heterogenization, and envision approaches that “embraces the variation” ^<u>Karp, Nicholson & Morgan</u>^. Heterogenization may be viewed as a way to simulate additional laboratories by varying the conditions within the original laboratory, a similar approach to their own approach of artificially increasing the variability in single-lab studies by adding the GxL interaction noise as measured in previous multi-lab studies ^<u>Benjamini, Kafkafi</u>^. However, Tucci suggested that rather than a systematic heterogenization it is worth investing effort in automated home-cage systems that can collect timestamps events in the cage, across multiple laboratories; therefore the strength of this approach would lie into the data sharing and a coordinated activity of analyses across mutant lines and across centers.

Surprisingly perhaps, it was agreed by almost all participants that extreme standardization is neither feasible nor helpful. Efforts had been made and should be made to develop better outcome measures and measurement systems. Given the pervasive nature of phenotypic plasticity resulting in GxE interaction, two directions were presented: Incorporate the interaction by statistical methods (see next section), or try to incorporate environmental variation in the experimental design rather than trying to spirit variation away through standardization.

## 8. Replicability across laboratories: can it be ensured?

The issue of replicability across laboratories is one of the most critical in mouse phenotyping, because modern science does not normally accept experimental results that are idiosyncratic to a certain location, even if they are replicable within this location. This is why the results and conclusions of the Crabbe et al. (1999) report were disturbing for many researchers in the field and in other fields as well. As previously discussed in section 6 there is currently no consensus even over the proper criteria to evaluate replicability across laboratories. Studies on the subject are few because they typically require collaboration of several laboratories and institutions, although they are becoming more and more common, thanks to data sharing and community efforts (section 3).

Therefore, credible and practical methodological and statistical solutions to the issue are urgently needed. Several strategies were discussed in the meeting, including the following proposals.

Ideally, discoveries should be replicated in at least one additional laboratory. In the simplest case of testing the same hypotheses of difference between two genotypes, e.g., a knockout and its wildtype control, the criterion for a replicated discovery may be statistical significance (using, e.g., a 0.05 level t-test) in each of two laboratories. Such a criterion concurs with the old Royal Society rule, as well as with Ronald Fisher’s view (see section 1). Unfortunately, this criterion is not trivial anymore when looking at p-values from multiple phenotypic measures, as is typical for mouse phenotyping, due to the issue of multiple comparisons. That is, with enough hypotheses tested this way, some of them will be found “replicable” just by coincidence. Heller et al. (2014) therefore generalized the criterion to multiple comparisons situations. She proposed a novel statistics for this purpose, the “r-value”. In the simplest case above the r-value equals the larger of the p-values in the two labs, but when multiple hypotheses are tested in each lab, the r-value can be generalized to take care of the multiple comparisons issue, using the FDR. Reporting the r-values can thus give credibility to the replicability claims. The r-value based on FDR is far more powerful than the alternative family-wise error method, and can be proved to control the expected fraction of false replicability claims.

In practice, however, most phenotyping experiments are done in a single laboratory, and results from other laboratories are usually not immediately available. This raises an unavoidable question: what should researchers do about significant discoveries in their own laboratory? How would they know if they are likely to replicate in other laboratories? Should they publish the discovery, or seek first to validate it in additional laboratories? And how would other researchers know if they are likely to replicate the effect in their own laboratory? All solutions discussed at this meeting have the effect of increasing the standard error of effect size, and many exciting findings that depend on exceeding the standard p < 0.05 threshold will not survive this simple test. A practical solution to these questions ^<u>Benjamini,</u> <u>Kafkafi,</u> Jaljule^ employs an extension of the random lab model (section 6), termed “GxL-adjustment”. The method enables researchers to apply the random lab model to phenotyping results in their own lab, providing a previous estimation of the interaction is available. The genotypic effect in the single laboratory is compared, as in the usual t-test, to the within-group variation, but this time “adjusted” by the addition of the multi-lab interaction variation. This addition of the interaction, as in the application of the random lab model to a multi-lab analysis, raises the benchmark for showing a significant genotype effect, ensuring that only effects that are likely to replicate in other laboratories will be significant. GxL-adjustment can be demonstrated to decrease the proportion of false discoveries that are not really replicable to the range of the traditional 0.05, for a price of modest reduction in the statistical power ^Jaljule^.

Several important insights can be gained from the Random Lab Model and from GxL-adjustment <u>^Benjamini^</u>. First, the size of the interaction variability sets its own limit for detection of replicable results. Thus, increasing the number of the animals within a single lab has only a limited benefit for the ability to replicate the results, since it does not affect the interaction with the laboratory. For the same reason, decreasing the individual animal noise has a limited benefit for replicability.

A phenotypic measure with smaller interaction is therefore “better” in the specific sense that it is more powerful to detect true replicable results, but not necessarily in other contexts. Consequently, we should search for phenotypic measures having smaller interaction, but keep in mind that replicability is a still property of the result, not of a phenotypic measure. That is, true replicable genotype differences can be apparent even over a large interaction, if they are large enough, and true replicable genotype differences that are small will be difficult to replicate even over a smaller interaction.

An extensive effort of standardization, as reported by Crabbe et al. (1999), might reduce individual noise, yet fail to eliminate unknown and unavoidable interaction sources. If individual noise is decreased but the interaction remains the same, the usual ANOVA model (with fixed lab effects) will paradoxically detect more significant interaction terms, giving a false impression of reduced replicability <u>^Kafkafi^</u>. The Random Lab Model in the same situation will give the correct impression: replicability (as seen in the number of significant genotype differences) will in fact improve, but only to a point. Beyond this point, further improvement of replicability must be achieved by decreasing the interaction.

The Random Lab Model does set a higher level for showing replicability. This is not necessarily a drawback, in the sense that it is a way to weed out non-replicable differences (Fonio et al., 2012). It is an important incentive to invest time and effort in reducing interaction. The interaction can be methodically reduced by improving analysis methods, e.g., robust smoothing (Benjamini et al., 2010). However, while interaction variability should be reduced, it will never be completely eliminated (much like the individual animal noise) and therefore should never be ignored. Unknown sources of interaction are unavoidable <u>^Würbel & Richter^</u>.

How can the interaction with the laboratory be estimated? One possibility is using as a surrogate the variability across time within a single laboratory <u>^Karp^</u> or “heterogenization” – injecting controlled variation through conditions within a single lab ^<u>Würbel & Richter</u>^. However, controlled heterogenization uses effects we know about, while true interaction with laboratory might involve factors we are not aware of at all. One proposal is to make use of multi-lab data from large and evolving phenotyping databases ^<u>Benjamini,</u> <u>Kafkafi,</u> Jaljule^, such as the Mouse Phenome Database (MPD) and the databases of the International Mouse Phenotyping Consortium (IMPC). Such database can calculate the interaction and publish it for use by scientists running phenotyping experiments in their own laboratories. This has to be repeated for each phenotypic measure separately. A website was demonstrated ^<u>Benjamini</u>^ in which any researcher conducting a phenotyping experiment can upload their results, get an updated estimate of the interaction for the relevant phenotypic measure, perform a GxL-adjustment and get an adjusted p-value. The researcher is then given an option to leave their data in the database, thus enriching it and providing a better estimate, based on more laboratories, for future users.

Ultimately, the replicability of a phenotyping discovery can be guaranteed only by testing it in additional laboratories. Even in these simple cases, ways to quantify replicability, such as the “r-value”, are still in need of development and acceptance by the scientific community. In cases when testing was performed in a single lab only, it may still be possible to enhance replicability, or estimate its prospects in a better way. Several directions were proposed: heterogenizing the experimental setup, splitting the duration of the experiments to different time units, and using external estimates of the interaction from phenotyping database. All these may serve to get more realistic estimates of the replicability standard deviation, and better (not too optimistic) estimates of the relevant standard errors.

## 9. Replicability and validity: what is the relation between them?

Several speakers in the meeting <u>^Chesler, Crabbe, Crusio, Gerlai, Williams, Würbel & Richter^</u> stressed the importance of validity of phenotyping results, and its probable close connection with replicability. Additional speakers <u>^Golani, Gomez-Marin, Pfaff, Tucci^</u> used different terms, yet they share the view that the issue of replicability may be a byproduct of a more fundamental methodological problem with behavioral phenotyping. There was no clear consensus over the practical meaning of the term “validity”, neither about how to ensure it, since there are several different kinds of validity, and different things often seemed valid to different speakers. However, generally all these speakers seem to share some dissatisfaction with the current credibility of phenotyping, especially behavioral phenotyping. As one of them noted: “What do we measure? Most of the time we have no clue.” <u>^Crusio^</u>. These speakers also seem to share the hope that, once phenotypes are properly validated, the issue of replicability will turn out to be considerably less grave as well.

Three kinds of validity are commonly employed in the discussion of animal models <u>^Chesler^</u>. It is expected that behavioral phenotypes that are closely related, even across different species such as mice and humans, should produce similar behaviors (“face validity”), should be influenced by homologous genes and CNS mechanisms (“construct validity”), and should respond in similar ways to similar experimental manipulations (“predictive validity”, e.g., Schellinck et al., 2010). Face validity is a problematic concept. Biological homology is a fundamental issue hardly attended to in behavioral phenotyping (Gomez-Marin et al., 2016). What we should regard as homologous behavior across species is not a simple question to answer using ad-hoc constructs. They require careful research of movement and behavior in species from different phyla, as remote as mice, humans and fruit flies. Once the common frame of reference and common origins used by organisms are identified, homology may be apparent in the invariable temporal and spatial order of movement components. This is similar to the way that homologous bones, despite dramatic variation in their shape and function across different vertebrate orders and classes, can readily be identified by their common relative position in the skeleton.

Viewed through these concepts, validity of measures may be obvious as it is in comparative anatomy, and even across mice, humans and flies.

Generalizability can be viewed as a sign of external validity <u>^Würbel & Richter^</u>. Replicability across laboratories is the least requirement. Replicability of behavioral phenotyps over particular housing and testing conditions are more valid. Replicability across more general conditions, such as different tests, mouse strains and even species can again be viewed as generalizability at different levels. Several other speakers seem to agree implicitly with this explicit assertion.

This view also appears similar to what is termed associative validity and dissociative validity <u>^Chesler^</u>. That is, the magnitude, frequency or quality of related phenotypic measures should be correlated, even across studies, genotypes and species, while unrelated phenotypic measures should be uncorrelated. A similar point was made by Crusio, who asked “what is it that we want to replicate?” <u>^Crusio^</u>. One view is that replicating results is very important practically, but replicating inferences – constructs such as “anxious” or “depressed” – is more important scientifically. In the example mentioned in section 5, the forced swimming test and the tail suspension test are often assumed to measure the same construct termed “behavioral despair”, yet this apparent face validity is not corroborated by predictive validity – mice that were treated with random stress procedure frequently scored the opposite in these two tests. This implies that validation and cross-validation of phenotyping assays are urgently needed <u>^Crusio^</u>.

GeneNetwork.org correlates different phenotypic assays and Mapping QTLs in the BXD recombinant inbred lines <u>^Williams^</u>. A high correlation between two phenotypic measures across many strains means suggests that these phenotypes measure a similar construct, even when they originate in different studies and different species, and measured in different tests and different conditions. Such correlated phenotypic measures may be viewed as different ways to measure the same trait that has been essentially replicated.

Moreover, if both phenotypic measures reveal the same strong QTL, the correlation implies a similar causal connection, since the central dogma assures us that it is the genotype that determines the phenotype rather than the other way around, and thus, construct validity in the genetic level is often gained as well.

A strategy of integrative bioinformatics was suggested as a way to discover validated and replicable relations among a variety of phenotypes through the shared association to
common genomic features <u>^Chesler^</u>. In a demonstration of this strategy, GeneWeaver (see section 3) was used to study the relationship between alcohol preference and alcohol withdrawal in gene sets from multiple publications and public data resources, including mouse QTL and humans (Bubier et al., 2014). Combinatorial integration of these gene sets revealed a single QTL positional candidate gene common to both preference and withdrawal. This QTL seems to have a replicable phenotypic effect – it was confirmed by a validation study in knockout mice mutated for this locus. Since discoveries in this strategy can be based on multiple studies across laboratories, species, and phenotyping protocols, they have a better chance to be replicable, generalizable, and translatable. However, the complex integration of multiple data sets in this strategy makes it difficult to construct statistical models for estimating how much the replicability may be improved.

A similar approach ^<u>Golani, Gomez-Marin</u>^ explicitly searches for measures of behavior that are more replicable across laboratories, generalized across conditions and test, and translatable across species. Both approaches integrate the outcomes of diverse experiments under varying conditions and test paradigms. The approaches are different in that the first combines existing phenotypic measures in various ways, while the second frequently redefine and formulate novel phenotypic measures by working directly with the low level movement data.

It therefore appears that most speakers agree about the importance of validity in behavioral phenotyping, but define it in different ways and employ different strategies to improve it. Some strategies are more “bottom-up”, searching for phenotypes that are already valid and replicable when measured in the individual animal, while other strategies are more “top-bottom”, hoping to construct valid and replicable inferences by integrating many of the current problematic phenotyping measures.

Many speakers consider validity, defined in various kinds, forms and levels, to be extremely important and worthy of addressing regardless of the replicability across laboratories issue. The current attention given to the methodological issue of replicability across laboratories may also help the more general issue of generalizability and validity.

**Overall Summary**

Modern science is reviewing its own problems of credibility, searching for causes and ways to address them. The original concepts of experimental science, emphasizing replicability and reproducibility, are still correct in spirit, but experimental, computational, and statistical methodologies require updating to cope with the increasing size and complexity of current research. Pre-clinical research and mouse phenotyping are similar in this regard to many other fields of experimental science. However, they enjoy special technical advantages of their own, such as the ability to standardize genomes and manipulate them in a precise manner, and also encounter special challenges, such as a potential interaction between genotype and environment, and the difficulties in defining and measuring behavioral phenotypes. Any proposed solutions, therefore, should likely be tailored to the particularities of the field. Phenotypic databases, community efforts and other methods of data sharing play important roles, as they can be employed to efficiently assess the severity of the issue, as well as the performances of different proposed solutions.

Correct handling of the genotype-environment interaction (GxE) is a key to proper methodology, and depends on the context and objective of the research. GxE can typically be understood and decreased through careful observation of the animals, redesigning the rearing and testing conditions to eliminate adverse effects of irrelevant confounding factors. Especially when testing a hypothesis of a genotype effect, ignoring or mishandling the relevant form of GxE is likely to result in replicability issues and other severe methodological issues. Extreme standardization of rearing and testing conditions is probably not by itself feasible or helpful as a strategy to eliminate GxE, and might limit the generalizability of any conclusions.

Ultimately, the replicability and generalizability of any phenotyping discovery can be guaranteed only by replicating it across additional studies, laboratories and experimental conditions. Even in situations when such replicating studies are available, there is no single well-established method to quantify the replicability of the results, but large and consistent genotype effect sizes can usually be agreed upon as replicable. In the more common situation where results are available from only one study, it may still be possible to enhance replicability, or estimate its prospects in a better way. Several directions were proposed and discussed at the meeting, including heterogenizing the experimental setup, splitting the duration of the experiments to different time units, and employing external estimates of the interaction from phenotyping database.

Linked with the issues of replicability are those relating to how phenotypes are defined and measured, especially for behavioral phenotypes. Regardless of the problems with replicating across laboratories, issues of generalizability and validity remain worth addressing. Insight and solutions resulting from attention given to the methodological issue of replicability may directly help with generalizability, and also help in addressing the more general issue of validity, by freeing investigators from rigid reliance on standardization, and rather promoting approaches to generalizable and replicable science. These observations first emerged from behavioral characterization of model organisms bear on other areas of biological inquiry and experimental science in general.

## Meeting Lectures

In the body of the paper, names in Upper case denote link to lecture videos of the participants in the meeting. The full lecture names and web addresses are given in the list below:

## Footnotes

2 <u>^Benjamini^</u> Yoav Benjamini: “Assessing Replicability in One Own's Lab: A Community Effort” <u>https://www.youtube.com/watch?v=y-Q4GWkJicE&list=PLNiWLB_wsOg74GlfLNyAcTo-TshyAcdLP&index=15</u>

3 <u>^Chesler^</u> Elissa J. Chesler: “Finding Consilience in Genetic and Genomic Analyses of Behavior” <u>https://www.youtube.com/watch?v=f9TdNXipRPQ&list=PLNiWLB_wsOg74GlfLNyAcTo-TshyAcdLP&index=20</u>

4 <u>^Crabbe^</u> John C. Crabbe: “Managing Sources of E to Address GXE” <u>https://www.youtube.com/watch?v=A7R2iZfjydA&list=PLNiWLB_wsOg74GlfLNyAcTo-TshyAcdLP&index=4</u>

5 <u>^Crusio^</u> Wim E. Crusio: “What do We Want to Reproduce?” <u>https://www.youtube.com/watch?v=3ZuEZw8mDjY&index=6&list=PLNiWLB_wsOg74GlfLNyAcTo-TshyAcdLP</u>

6 <u>^Eilam^</u> David Eilam: “Variability: A By-Product Noise or an Essential Component of Motor Behavior” <u>https://www.youtube.com/watch?v=ADgVHDFZpUg&list=PLNiWLB_wsOg74GlfLNyAcTo-TshyAcdLP&index=7</u>

7 <u>^Heller^</u> Ruth Heller: “Assessing Replicability Across Laboratories: The R-Value” <u>https://www.youtube.com/watch?v=b2uGOp9yLMw&list=PLNiWLB_wsOg74GlfLNyAcTo-TshyAcdLP&index=22</u>

8 <u>^Gerlai^</u> Robert Gerlai: “Behavioral Phenotyping: The Double Edged Sword” <u>https://www.youtube.com/watch?v=pRrNz0PTKc4&list=PLNiWLB_wsOg74GlfLNyAcTo-TshyAcdLP&index=9</u>

9 <u>^Golani^</u> Ilan Golani: “Replicability as a Virtue” <u>https://www.youtube.com/watch?v=KiRSxpA8qZY&index=10&list=PLNiWLB_wsOg74GlfLNyAcTo-TshyAcdLP&spfreload=10</u>

10 <u>^Gomez-Marin^</u> Alex Gomez-Marin: “Toward a Behavioral Homology Between Insects and Rodents” <u>https://www.youtube.com/watch?v=Hu8PwdKap3k&list=PLNiWLB_wsOg74GlfLNyAcTo-TshyAcdLP&index=25&spfreload=10</u>

11 <u>^Iraqi^</u> Fuad Iraqi: “TAU Collaborative Cross Mice a Powerful GRP for Dissection Host Susceptibility to Diseases” <u>https://www.youtube.com/watch?v=NV62o7Ubrfg&list=PLNiWLB_wsOg74GlfLNyAcTo-TshyAcdLP&index=21</u>

12 <u>^Karp^</u> Natasha A. Karp: “Simulation Studies to Investigate Workflow and its Impact on Reliability of Phenodeviant Calls: <u>https://www.youtube.com/watch?v=N5wTrgXvsAY&index=17&list=PLNiWLB_wsOg74GlfLNyAcTo-TshyAcdLP</u>

13 <u>^Kafkafi^</u> Neri Kafkafi: “The Random Lab Model for Assessing the Replicability of Phenotyping Results Across Laboratories” <u>https://www.youtube.com/watch?v=XsQawSBA6Vc&list=PLNiWLB_wsOg74GlfLNyAcTo-TshyAcdLP&index=12</u>

14 <u>^Nicholson & Morgan^</u> George Nicholson & Hugh Morgan: “The Empirical Reproducibility of High-Throughput Mouse Phenotyping” <u>https://www.youtube.com/watch?v=KzrrP6_F8r8&index=18&list=PLNiWLB_wsOg74GlfLNyAcTo-TshyAcdLP</u>

15 <u>^Pfaff^</u> Donald W. Pfaff: “Application of Strict Methodology and Applied Mathematical Statistics to Mouse Behavioral Data” <u>https://www.youtube.com/watch?v=hB0FnO9evbY&index=8&list=PLNiWLB_wsOg74GlfLNyAcTo-TshyAcdLP</u>

16 <u>^Stark^</u> Philip B. Stark: “Preproducibility for Research, Teaching, Collaboration, and Publishing” <u>https://www.youtube.com/watch?v=wHryMtEBkB4&list=PLNiWLB_wsOg74GlfLNyAcTo-TshyAcdLP&index=13</u>

17 <u>^Stiedl^</u> Oliver Stiedl: “Individuality of Avoidance Behavior of Mice in an Automated Home Cage Environment” <u>https://www.youtube.com/watch?v=gUIgRW5luZY&list=PLNiWLB_wsOg74GlfLNyAcTo-TshyAcdLP&index=24</u>

18 <u>^Stodden^</u> Victoria Stodden: “Computational and Statistical Reproducibility” <u>https://www.youtube.com/watch?v=GzrOcqz8TVY&index=14&list=PLNiWLB_wsOg74GlfLNyAcTo-TshyAcdLP</u>

19 <u>^Tucci^</u> Valter Tucci: “Phenotyping Behaviour Across Laboratories and Across Mouse Strains” <u>https://www.youtube.com/watch?v=iTlsFaj62oQ&index=22&list=PLNiWLB_wsOg74GlfLNyAcTo-TshyAcdLP</u>

20 <u>^Valdar & Tarantino^</u> William Valdar & Lisa M. Tarantino: “The Effect of Genetic Background on Behavioral Variability: Implications for Replicability?” <u>https://www.youtube.com/watch?v=63sgLO4Hd04&list=PLNiWLB_wsOg74GlfLNyAcTo-TshyAcdLP&index=5</u>

21 <u>^Williams^</u> Robert W. Williams: “Data Rescue for Replication: Finding, Annotating, and Reusing Data for the BXD Mouse Cohort” <u>https://www.youtube.com/watch?v=goocssSA33g&index=19&list=PLNiWLB_wsOg74GlfLNyAcTo-TshyAcdLP</u>

22 <u>^Würbel & Richter^</u> Hanno Würbel & S. Helene Richter: “On Standardization and Other Fallacies in Animal Phenotyping” <u>https://www.youtube.com/watch?v=tfW35740q3k&index=11&list=PLNiWLB_wsOg74GlfLNyAcTo-TshyAcdLP</u>

## References

1. Agassi, J., 2013. “The very idea of modern science: Francis Bacon and Robert Boyle”. Boston Studies in the Philosophy and History of Science 298, Springer Science+Busness Media, Dordrecht. doi:10.1007/978-94-007-5351-8_1

2. Alberts, B., Cicerone, R. J., Fienberg, S. E., Kamb, A., McNutt, M., Nerem, R. M., Schekman, R., Shiffrin, R., Stodden, V., Suresh, S., Zuber, M. T., Pope, B. K., Jamieson, K. H., 2015. Scientific integrity: Self-correction in science at work. Science 348(6242), 1420–1422. doi: 10.1126/science.aab3847.

3. Anderson, P. W. 1972. More is different, Science, 177(4047), 393–396.

4. Anisman, H., Zaharia, M. D., Meaney, M. J., Merali Z., 1998. Do early-life events permanently alter behavioral and hormonal responses to stressors? Int. J. Dev. Neurosci. 16(3-4),149–164. doi:10.1016/S0736-5748(98)00025-2.

5. de Angelis M. H. et al., 2015. Analysis of mammalian gene function through broad-based phenotypic screens across a consortium of mouse clinics. Nat. Genet. 47(9):969–978. doi: 10.1038/ng.3360.

6. Asendorpf, A. G., et al., 2013. Recommendations for Increasing Replicability in Psychology. European Journal of Personality, Eur. J. Pers. 27, 108–119.

7. Baggerly, K. A., Coombes, K. R. 2009. Deriving chemosensitivity from cell lines: Forensic bioinformatics and reproducible research in high-throughput biology. Ann. Appl. Stat 3(4), 1309–1334.

8. Baker, E. J, Jay, J. J., Bubier, J. A., Langston, M. A., Chesler, E. J., 2012. GeneWeaver: a web-based system for integrative functional genomics. Nucleic Acids Res. D1067–1076. doi: 10.1093/nar/gkr968.

9. Beckers, J., Wurst, W., & de Angelis, M. H., 2009. Towards better mouse models: enhanced genotypes, systemic phenotyping and envirotype modelling. Nat. Rev. Genet. 10(6), 371–380. doi:10.1038/nrg2578.

10. Benjamini, Y., Hochberg, Y. 1995. Controlling the false discovery rate: a practical and powerful approach to multiple testing. J. Roy. Stat. Soc. B (Methodological), 1995, 289–300.

11. Benjamini, Y., Drai, D., Elmer, G., Kafkafi, N., Golani, I., 2001. Controlling the false discovery rate in behavior genetics research. Behav. Brain Res. 125(1-2), 279–284.

12. Benjamini, Y., Lipkind, D., Horev, G., Fonio, E., Kafkafi, N., Golani, I., 2010. Ten ways to improve the quality of descriptions of whole-animal movement. Neurosci. Biobehav. Rev. 34(8), 1351–1365. doi:10.1016/j.neubiorev.2010.04.004

13. Benjamini, Y., Hechtlinger, Y., 2013. Discussion: an estimate of the science-wise false discovery rate and applications to top medical journals by Jager and Leek. Biostatistics, 15(1), 13–16; discussion 39–45. doi: 10.1093/biostatistics/kxt032.

14. Brown, R. E., Stanford, L., Schellinck, H. M., 2000. Developing standardized behavioral tests for knockout and mutant mice. ILAR Journal, 41(3), 163–174.

15. 15.

16. Bubier, J. A., Jay, J. J., Baker, C. L., Bergeson, S. E., Ohno, H., Metten, P., Crabbe, J. C., Chesler, E. J., 2014. Identification of a QTL in Mus musculus for Alcohol Preference, Withdrawal, and Ap3m2 Expression Using Integrative Functional Genomics and Precision Genetics. Genetics 197(4), 1377–1393. doi:10.1534/genetics.114.166165

17. Chesler, E. J., Wilson, S. G., Lariviere, W. R., Rodriguez-Zas, S. L., Mogil, J. S., 2002. Identification and ranking of genetic and laboratory environment factors influencing a behavioral trait, thermal nociception, via computational analysis of a large data archive, Neurosci. Biobehav. Rev. doi:10.1016/S0149-7634(02)00103-3.

18. Chesler, E. J., Miller, D. R., Branstetter, L.R., Galloway, L. D., Jackson, B. L., Philip, V. M., Voy, B. H., Culiat, C. T., Threadgill, D. W., Williams, R. W., Churchill, G. A., Johnson, D. K., Manly, K. F. 2008. The Collaborative Cross at Oak Ridge National Laboratory: developing a powerful resource for systems genetics. Mamm Genome 19(6), 382–389. doi:10.1007/s00335-008-9135-8

19. Collaborative Cross Consortium, 2004. The Collaborative Cross, a community resource for the genetic analysis of complex traits. Nat. Genet. 36(11), 1133–1137.

20. Collaborative Cross Consortium, 2012. The genome architecture of the Collaborative Cross mouse genetic reference population. Genetics 190(2), 389–401. doi: 10.1534/genetics.111.132639.

21. Collins, F. S., Rossant, J., & Wurst, W., 2007. A mouse for all reasons. Cell 128(1), 9–13. http://dx.doi.org/10.1016/j.cell.2006.12.018.

22. Collins, F. S., Tabak, L. A., 2014. Policy: NIH plans to enhance reproducibility. Nature 505, 612–613. doi:10.1038/505612a.

23. Complex Trait Consortium. 2003. The nature and identification of quantitative trait loci: a community’s view. Nat Rev Genet, 4(11), 911.

24. Crabbe, J. C., Wahlsten, D., Dudek, B. C., 1999. Genetics of mouse behavior: interactions with laboratory environment, Science. 284(5420):1670–1672.

25. Crusio, W. E., 2006. Inheritance of behavioral and neuroanatomical phenotypical variance: hybrid mice are not always more stable than inbreds. Behav Genet, 36(5), 723–731.

26. Diggle, P. J., Zeger, S. L., 2010. Embracing the concept of reproducible research. Biostatistics 11(3), 375. doi: 10.1093/biostatistics/kxq029.

27. Eilam, D., 2014. The cognitive roles of behavioral variability: idiosyncratic acts as the foundation of identity and as transitional, preparatory, and confirmatory phases. Neurosci. Biobehav. Rev. 49:55–70. doi: 10.1016/j.neubiorev.11.023. Epub 2014 Dec 13.

28. Errington, T. M., Iorns, E., Gunn, W., Tan, F. E., Lomax, J., Nosek, B. A., 2014. An open investigation of the reproducibility of cancer biology research. Elife, 3, e04333.

29. Fisher, R. A., 1935. The Design of Experiments, Oliver & Boyd. Oxford, England, p. 14.

30. Fonio, E., Benjamini, Y., Golani, I., 2009. Freedom of movement and the stability of its unfolding in free exploration of mice. Proc. Natl. Acad. Sci. USA. 106(50), 21335–21340. doi: 10.1073/pnas.0812513106.

31. Fonio, E., Golani, I., Benjamini, Y., 2012. Measuring behavior of animal models: faults and remedies. Nat. Meth. 9(12), 1167.

32. Gerlai, R. 2002. Phenomics: fiction or the future? Trends Neurosci. 25(10), 506–509 doi:10.1016/S0166-2236(02)02250-6.

33. Gilbert, D. T., King, G, Pettigrew, S, Wilson, T. D., 2016. Comment on “Estimating the reproducibility of psychological science”. Science 351(6277), 1037, DOI: 10.1126/science.aad7243.

34. Gomez-Marin, A., Paton, J. J Kampff, A. Costa, R, R., Zachary, M., Mainen, M., 2014. Big behavioral data: psychology, ethology and the foundations of neuroscience. Nat. Neurosci. 17, 1455–1462.

35. Gomez-Marin, A., Oron, E., Gakamsky, A., Valente, D., Benjamini, Y., Golani, I., 2016. Generative rules of Drosophila locomotor behavior as a candidate homology across phyla. Sci. Rep. 6, 27555.

36. Goodman, S. N., Fanelli, D., Ioannidis, J. P., 2016. What does research reproducibility mean? Sci. Transl. Med. 8(341), 341ps12–341ps12.

37. Grubb, S. C., Bult, C. J., Bogue, M. A. 2014 Mouse phenome database. Nucleic Acids Res. Jan;42(Database issue):D825–34. doi: 10.1093/nar/gkt1159.

38. Hager, T., Jansen, R. F., Pieneman, A. W., Manivannan, S. N., Golani, I., van der Sluis, S., Smit AB, Verhage, M., Stiedl O., 2014. Display of individuality in avoidance behavior and risk assessment of inbred mice. Front Behav. Neurosci. 8, 314.

39. Heller, R., Bogomolov, M., Benjamini, Y., 2014. Deciding whether follow-up studies have replicated findings in a preliminary large-scale “omics’ study”. Proc. Natl. Acad. Sci. USA 111(46), 16262–16267.

40. Homberg, J. R et al. 2016. Understanding autism and other neurodevelopmental disorders through experimental translational neurobehavioral models. Neurosci. Biobehav. Rev. 65, 292–312.

41. Houtkooper, R. H., Mouchiroud, L., Ryu, D., Moullan, N., Katsyuba, E., Knott, G., Williams, R. W., Auwerx, J., 2013. Mitonuclear protein imbalance as a conserved longevity mechanism. Nature. 497(7450), 451–457. doi:10.1038/nature12188.

42. Ioannidis, J., 2005. Why most published research findings are false. PLoS Med. 2(8):e124.

43. Iraqi, F. A., Mahajne, M., Salaymeh A, Sandovsky H, Tayem H, Vered K, et al. 2012. The genome architecture of the Collaborative Cross mouse genetic reference population. Genetics; 190, 389–401.

44. Iraqi, F. A., Churchill, G., & Mott, R., 2008. The Collaborative Cross, developing a resource for mammalian systems genetics: a status report of the Wellcome Trust cohort. Mammalian Genome, 19(6), 379–381.

45. Jager, L. R., Leek, J. T. 2014. An estimate of the science-wise false discovery rate and application to the top medical literature. Biostatistics, 15(1), 1–12. doi:10.1093/biostatistics/kxt007.

46. Kafkafi, N., Benjamini, Y., Sakov, A., Elmer, G. I., Golani, I., 2005. Genotype-environment interactions in mouse behavior: a way out of the problem. Proc. Natl. Acad. Sci. USA. 102(12), 4619–4624.

47. Karp, N. A., Speak, A. O., White, J. K., Adams, D. J., de Angelis, M. H., Hérault, Y., Mott, R. F., 2014. Impact of Temporal Variation on Design and Analysis of Mouse Knockout Phenotyping Studies. 9(10), e111239. PLoS One, doi:10.1371/journal.pone.0111239

48. Keeley, P. W., Zhou, C., Lu, L., Williams, R. W., Melmed, S., Reese, B. E., 2014 Pituitary tumor-transforming gene 1 regulates the patterning of retinal mosaics. Proc. Natl. Acad. Sci. USA. 111(25), 9295–9300. doi: 10.1073/pnas.1323543111.

49. Kennet, R. S., Shmueli, G., 2015. Clarifying the terminology that describes scientific reproducibility. Nat. Meth. 12, 699. doi:10.1038/nmeth.3489.

50. Koscielny, G., et al. 2014. The International Mouse Phenotyping Consortium Web Portal, a unified point of access for knockout mice and related phenotyping data. Nucleic Acids Res. 42 (D1), D802–D809. doi: 10.1093/nar/gkt977.

51. Koutnikova, H., et al., 2009. Identification of the UBP1 locus as a critical blood pressure determinant using a combination of mouse and human genetics. PLoS Genet. 5(8):e1000591. doi: 10.1371/journal.pgen.1000591.

52. Landis, S. C., et al., 2012. A call for transparent reporting to optimize the predictive value of preclinical research. Nature 490, 187–191. doi:10.1038/nature11556.

53. Lander, E., Kruglyak, L. 1995. Genetic dissection of complex traits: guidelines for interpreting and reporting linkage results. Nat. Genet. 11(3), 241–247. doi:10.1038/ng1195-241.

54. Leek, J. T., Peng, R. D., 2015. Opinion: Reproducible research can still be wrong: Adopting a prevention approach. Proc. Natl. Acad. Sci. USA 112 (6), 1645–1646 (2015) doi: 10.1073/pnas.1421412111.

55. Lewontin, R. C. (1974). Annotation: the analysis of variance and the analysis of causes. Am. J. Human Genet. 26(3), 400.

56. McNutt, M., 2014. Reproducibility. Science 343 (6168), 229. doi:10.1126/science.1250475.

57. Morgan, H. et al. 2009. EuroPhenome: a repository for high-throughput mouse phenotyping data. Nucleic Acids Res. 38 (Issue suppl 1), D577–D585.

58. Morahan, G., Balmer, L., Monley, D. 2008 Establishment of “The Gene Mine”: a resource for rapid identification of complex trait genes. Mamm Genome. 19(6), 390–393. doi: 10.1007/s00335-008-9134-9.

59. Nature Editorial 2013, Announcement: Reducing our irreproducibility. Nature 496, 398. doi:10.1038/496398a.

60. Open Science Collaboration (2015) Estimating the reproducibility of psychological science. Science, 349(6251), aac4716 doi: org/10.1126/science.aac4716.

61. Pigliucci, M., 2001. Phenotypic plasticity: beyond nature and nurture. JHU Press.

62. Pollin, R., 2014. Public debt, GDP growth, and austerity: why Reinhart and Rogoff are wrong, LSE American Politics and Policy, - eprints.lse.ac.uk

63. Potti, A. et al. 2006 (retracted) A genomic strategy to refine prognosis in early-stage non-small-cell lung cancer. N. Engl. J. Med. 355, 570.

64. Peng, R. 2011. Reproducible Research in Computational Science, Science 334(6060), 1226–1227. doi: 10.1126/science.1213847

65. Peng, R., 2015. The reproducibility crisis in science: A statistical counterattack. Significance 12(3), 30–32.

66. Pfaff, D. W., 2001. Precision in mouse behavior genetics. Proc. Natl. Acad. Sci. USA. 98(11), 5957–5960.

67. Proekt, A., Banavar, J. R., Maritan, A., Pfaff, D. W., 2012. Scale invariance in the dynamics of spontaneous behavior. Proc. Natl. Acad. Sci. USA. 109(26), 10564–10569. doi: 10.1073/pnas.1206894109.

68. Richter, S. H., Garner, J. P., Würbel, H., 2009. Environmental standardization: cure or cause of poor reproducibility in animal experiments? Nat. Meth., 6, 257–261

69. Richter, S. H., Auer, C., Kunert, J., Garner, J. P., Würbel, H. 2010. Systematic variation improves reproducibility of animal experiments. Nat. Meth. 7, 167–168.

70. Richter, S. H. et al. (2011) Effect of Population Heterogenization on the Reproducibility of Mouse Behavior: A Multi-Laboratory Study. PLoS One 6(1), e16461. doi: 10.1371/journal.pone.0016461

71. Savalei, V., Dunn, E., 2015. Is the call to abandon p-values the red herring of the replicability crisis? Frontiers in psychology, 6, 245. http://dx.doi.org/10.3389/fpsyg.2015.00245

72. Stodden, V., 2010, The scientific method in practice: Reproducibility in the computational sciences. PLoS One DOI: 10.1371/journal.pone.0067111

73. Stodden, V., Guo, P. Ma, Z. 2013, Toward Reproducible Computational Research: An Empirical Analysis of Data and Code Policy Adoption by Journals

74. Stodden V., 2013. Resolving Irreproducibility in Empirical and Computational Research. http://bulletin.imstat.org/2013/11/resolving-irreproducibility-in-empirical-and-computational-research/

75. Shapin, S., Schaffer, S., 1985, Leviathan and the air-pump. Princeton University Press, Princeton, NJ.

76. Soric, B. 1987. Statistical “discoveries” and effect-size estimation. J. Am. Stat. Ass. 84(406), 608–610.

77. J. P. Simmons, Nelson, L. D., Simonsohn, U., 2011. False-positive psychology undisclosed flexibility in data collection and analysis allows presenting anything as significant. Psychol. Sci. 0956797611417632. 10.1177/0956797611417632

78. Soergel DAW. Rampant software errors may undermine scientific results [version 2; referees: 2 approved]. F1000Research 2015, 3:303. doi:10.12688/f1000research.5930.2

79. https://www.nist.gov/news-events/news/2006/10/new-web-based-system-leads-better-more-timely-data.

80. Valdar, W., Solberg, L. C., Gauguier, D., Cookson, W. O., Rawlins, J. N., Mott, R, Flint, J., 2006. Genetic and environmental effects on complex traits in mice. Genetics. 174(2), 959–984.

81. Voelkl, B., and Würbel, H., 2016. Reproducibility Crisis: Are We Ignoring Reaction Norms? Trends Pharmacol. Sci., 37, 509–510.

82. Vul, E., Harris, C., Winkielman, P., Pashler, H. (2009) Puzzlingly High Correlations in fMRI Studies of Emotion, Personality, and Social Cognition. Perspect Psychol Sci, 4(3):274–90. doi: 10.1111/j.1745-6924.2009.01125.x.

83. Wang et al., 2016, Joint mouse-human phenome-wide association to test gene function and disease risk. Nat. Comm. 10464. doi: 10.1038/ncomms10464.

84. Williams, E. G., Auwerx, J. 2015. The Convergence of Systems and Reductionist Approaches in Complex Trait Analysis. Cell 162(1), 23–32. doi:10.1016/j.cell.2015.06.024.

85. Wolfer, D. P., Stagljar-Bozicevic, M., Errington, M. L., Lipp, H. P., 1998. Spatial Memory and Learning in Transgenic Mice: Fact or Artifact? Physiology, 13 (3), 118–123.

86. Wolfer, D. P. et al. 2004. Laboratory animal welfare: cage enrichment and mouse behaviour. Nature 432(7019), 821–822.

87. Würbel, H., 2000. Behaviour and the standardization fallacy. Nat. Genet. 26, 263. doi:10.1038/81541.

88. Wahlsten, D., 1990. Insensitivity of the analysis of variance to hereditary-environment interaction. Behav. Brain Sci. 13, 109–120.

89. Wahlsten, D., Rustay, N. R., Metten, P., & Crabbe, J. C., 2003. In search of a better mouse test. Trends Neurosci. 26(3), 132–136. doi:10.1016/S0166-2236(03)00033-X

90. Wahlsten, D., Bachmanov, A., Finn, D. A., Crabbe, J. C., 2006. Stability of inbred mouse strain differences in behavior and brain size between laboratories and across decades. Proc Nat Acad Sci USA 103, 16364–16369.

91. Welsh, C. E., Miller, D. R., Manly, K. F., Wang, J., McMillan, L., Morahan, G., et al. Status and access to the Collaborative Cross population. Mamm. Genome 23, 706–712.

